# Low genetic variation is associated with low mutation rate in the giant duckweed

**DOI:** 10.1101/381574

**Authors:** Shuqing Xu, Jessica Stapley, Saskia Gablenz, Justin Boyer, Klaus J. Appenroth, K. Sowjanya Sree, Jonathan Gershenzon, Alex Widmer, Meret Huber

## Abstract

Mutation rate and effective population size (*N_e_*) jointly determine intraspecific genetic diversity, but the role of mutation rate is often ignored. We investigate genetic diversity, spontaneous mutation rate and *N_e_* in the giant duckweed (*Spirodela polyrhiza*). Despite its large census population size, whole-genome sequencing of 68 globally sampled individuals revealed extremely low within-species genetic diversity. Assessed under natural conditions, the genome-wide spontaneous mutation rate is at least seven times lower than estimates made for other multicellular eukaryotes, whereas *N_e_* is large. These results demonstrate that low genetic diversity can be associated with large-*N_e_* species, where selection can reduce mutation rates to very low levels, and accurate estimates of mutation rate can help to explain seemingly counterintuitive patterns of genome-wide variation.

**One Sentence Summary:** The low-down on a tiny plant: extremely low genetic diversity in an aquatic plant is associated with its exceptionally low mutation rate.

## Main Text

Explaining within-species genetic diversity—measured as the level of intraspecific DNA sequence variation—is a major goal in evolutionary and conservation biology, as this diversity can influence how species cope with changing environments (*1, 2*). While intraspecific genetic diversity is known to vary widely among species, the underlying causes remain controversial (*3, 4*). According to population genetic theory, the population mutation parameter (*θ*) is determined by the product of the spontaneous neutral mutation rate (*μ*) and effective population size (*N_e_*), and in diploid species *θ*= 4 × *N_e_* × *μ* (*5*). In practice, the parameter *θ* is often estimated by the average pairwise nucleotide diversity (π) at putatively neutral sites (*6*). While the role of *N_e_* in explaining variation in genetic diversity among taxa has received much theoretical and empirical attention (*3, 4, 7*), the influence of variation in mutation rate and the interaction between *N_e_* and mutation rate remain largely unknown.

As most spontaneous mutations are deleterious, selection should favor lower mutation rates, but in small populations the efficacy of selection to lower the mutation rate is limited as genetic drift overrides the effect of natural selection. This ‘drift-barrier’ hypothesis can explain variation in mutation rates and the observed negative relationship between effective population size and mutation rate among species (*8*). However, one counter-intuitive prediction of the drift-barrier hypothesis is that populations with large *N_e_* may also have low genetic diversity if natural selection has driven mutation rates to very low levels. Whether this pattern is present in eukaryotes is unknown, largely due to the paucity of studies quantifying both genome-wide diversity and spontaneous mutation rates under natural conditions in organisms with different life histories and reproductive strategies.

To better understand the relationship between genetic diversity, mutation rate and *N_e_*, we independently obtained genome- and range-wide estimates of genetic diversity and mutation rate in the diploid freshwater plant *Spirodela polyrhiza* L. (Schleid.) (‘duckweed’; Lemnaceae). This species is one of the fastest growing angiosperms; under suitable growth conditions, it reproduces predominantly by asexual budding with a duplication rate of 2-3 days (*9, 10*). Consequently, *S. polyrhiza* often achieves extremely high census population sizes in nature as millions of individuals can be found in a single pond. However, previous studies using a limited number of genetic markers found low genetic diversity (*11, 12*).

To provide genome- and range-wide estimates of genetic diversity in *S. polyrhiza*, we resequenced the genomes of 68 genotypes representing the global distribution of the species, using Illumina short-read sequencing with 29X average coverage (Table S1). All sequence reads were aligned to the *S. polyrhiza* reference genome (*14*) using the BWA-MEM aligner and genetic variants were identified using GATK (*15*). In total, we found 996,115 biallelic and 7,880 multiallelic high-quality single nucleotide polymorphisms (SNPs) as well as 214,262 small indels. This represents on average one SNP per 145 bp in the *S. polyrhiza* genome, which is low compared to an average of 1 SNP per 23 bp in *Arabidopsis thaliana* when a comparable number of genotypes are sequenced (*16*). Among all biallelic SNPs, 14,191 nonsynonymous and 8,865 synonymous SNPs were found (Table S2 and External Dataset 1). The estimated *S. polyrhiza* range-wide pairwise nucleotide diversity at synonymous sites (π_s_) was 0.00086, which is among the lowest values reported for any multicellular eukaryote for which genome-wide genetic diversity has been estimated (Table S3) (*3*).

Population structure analysis based on genome-wide polymorphisms revealed four population clusters in *S. polyrhiza*, which are centered in four geographic regions: America, Europe, India and South East (SE) Asia (Figure 1). A few samples showed discrepancies between their geographic origin and population cluster assignment based on their genomic variation, likely due to either recent migrations of the duckweed associated with human activities or mis-labeling during long-term maintenance of the duckweed collections. The pairwise *F_st_*, an indicator of relative differentiation between populations, ranged from 0.35 to 0.79 (Table S4), suggesting distinct regional populations in *S. polyrhiza*. Between populations, the genome-wide nucleotide diversity from all sites ranged between 0.00067 (SE Asian versus European population) and 0.00013 (European versus American population). Within populations, π calculated from all sites ranged from 0.00018 (American population) to 0.00056 (SE Asian population) (Table S5). The extent of linkage disequilibrium (LD), measured as the average distance between variants when their correlation coefficient (*r*^2^) = 0.2, varied from 14.1 kb in the SE Asian population to 143.2 kb in the European population. The relatively slow decay of LD in *S. polyrhiza* may be attributed to its predominantly clonal reproduction. Comparing across populations, we observed much faster LD decay in the SE Asian population, suggesting more frequent (historical or ongoing) sexual reproduction in this region and/or higher *N_e_*. Together, these results establish that genome-wide nucleotide diversity in *S. polyrhiza* is extremely low and sexual reproduction might be more frequent in the SE Asian population.

**Fig. 1.**
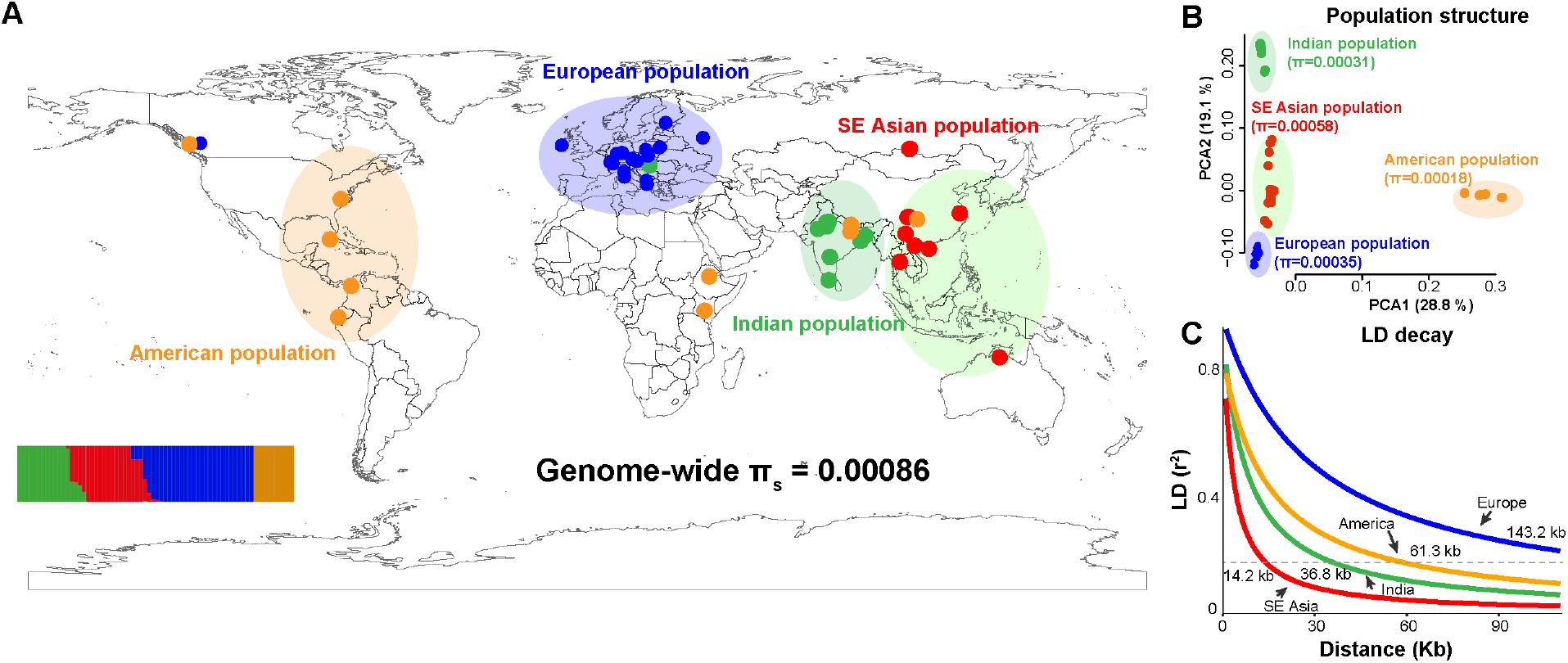
Genome-wide nucleotide diversity, population structure and linkage disequilibrium in *S. polyrhiza*. (**A**) Geographic distribution of the 68 sequenced samples, colored according to population structure. The insert at the lower left corner shows the results from the STRUCTURE analysis using genome-wide polymorphisms. Each colored line refers to an individual and the Y-axis refers to the likelihood. Genome wide π_s_ refers to average pairwise nucleotide diversity at synonymous sites. (**B**) Principal Coordinate Analysis (PCA) based on genome-wide nucleotide diversity data. Average pairwise nucleotide diversity (π) calculated from all sites is shown for each population. (**C**) Decay of linkage disequilibrium (LD) with physical distance in four populations. The dashed line indicates an LD value of *r*^2^ = 0.2, and the numbers refer to the mean pairwise distance between sites at *r*^2^ = 0.2.

To investigate if the observed low genomic diversity in *S. polyrhiza* can be explained by universally low mutation rate or, alternatively, low effective population size, we estimated the spontaneous mutation rate and used our estimates of mutation rate and genomic diversity to estimate effective population size. Mutation rates can be markedly affected by outdoor environmental stresses such as temperature fluctuations and ultraviolet (UV) light (*17-21*), conditions that prevail in the native habitats of *S. polyrhiza*. Consequently, we estimated the genomic mutation rate in indoor and outdoor mutation accumulation (MA) experiments, and manipulated UV light in the outdoor experiments to further assess the effect of environmental stresses (Figure S1 and External Dataset 2). Offspring of a single common ancestor were propagated as single descendants under these conditions for 20 generations (Figure S2), after which individual plants from five replicates per treatment were collected, and their genomes sequenced and compared to the ancestral genome. We obtained genome information for 16 individuals (including the common ancestor) with an average coverage of 28X (Table S6) and identified genetic variants in more than 79.7% of the *S. polyrhiza* genome (~126 Mb). Among the 15 offspring, four *de novo* mutations were identified and confirmed by Sanger sequencing. These mutations all originated from the outdoor MA experiments, and located in non-coding regions. One mutation (C:G->T:A) was found in a UV-shielded line and the other three mutations (two C:G->T:A and one C:G->A:T) were found in UV-exposed lines (Table 1). Further analysis that compared the heterozygous sites of maternal and offspring individuals suggested a low false-negative rate for our mutation identification pipeline (1.6 ± 0.7%). Given that the protein-coding region of the *S. polyrhiza* genome is 17.4 Mb, we estimate the number of mutations per generation in the entire protein-coding DNA of *S. polyrhiza* under natural, outdoor conditions to be 0.0042 ± 0.0038. As so few mutations were observed, we were unable to perform robust statistical analysis. However, the higher number of mutations found in outdoor samples and in the presence of UV light is consistent with the hypothesis that outdoor stresses increase the spontaneous mutation rate.

**Table 1.** Summary of the sequencing data and detected mutations. Each row shows the sample information and number of verified mutations. Effective sites are estimated as the total number of sites with sufficient coverage for finding *de novo* variants using our pipeline. The mutation rate was calculated as *μ* = (number of mutations / sum of effective sites) / number of generations.

| Sample ID | Treatment | # Mutations | Effective sites (Mb) | Average $\mu$ |
| --- | --- | --- | --- | --- |
| A | Indoor | 0 | 126.4 | <7.92E-11 |
| E | Indoor | 0 | 125.7 |  |
| I | Indoor | 0 | 126.0 |  |
| J | Indoor | 0 | 126.4 |  |
| N | Indoor | 0 | 125.9 |  |
| B | Outdoor-noUV | 0 | 126.1 | 7.92E-11 |
| G | Outdoor-noUV | 1 | 126.3 |  |
| K | Outdoor-noUV | 0 | 124.2 |  |
| O | Outdoor-noUV | 0 | 125.9 |  |
| P | Outdoor-noUV | 0 | 126.4 |  |
| C | Outdoor-UV | 1 | 125.6 | 2.38E-10 |
| D | Outdoor-UV | 1 | 126.3 |  |
| L | Outdoor-UV | 1 | 126.3 |  |
| M | Outdoor-UV | 0 | 126.3 |  |
| Q | Outdoor-UV | 0 | 126.0 |  |

The genome-wide mutation rate in *S. polyrhiza* is within the range of mutation rates reported for unicellular eukaryotes and Eubacteria, but is more than seven times lower than the reported rates for multicellular eukaryotes (Figure 2). This estimated seven-fold difference between *S. polyrhiza* and other multicellular eukaryotes is a conservative estimate, as all MA experiments in other organisms were performed under controlled indoor conditions, which likely resulted in lower mutation rate estimates. Based on these independent estimates of genetic diversity and mutation rate, we can estimate *N_e_* in *S. polyrhiza*. Assuming that mutation rates during the clonal and sexual reproduction phases of *S. polyrhiza* are equal, the estimated effective population size of *S. polyrhiza* is 9.0 × 10^5^, which is among the highest estimates for multicellular eukaryotes (Table S3).

**Fig. 2.**
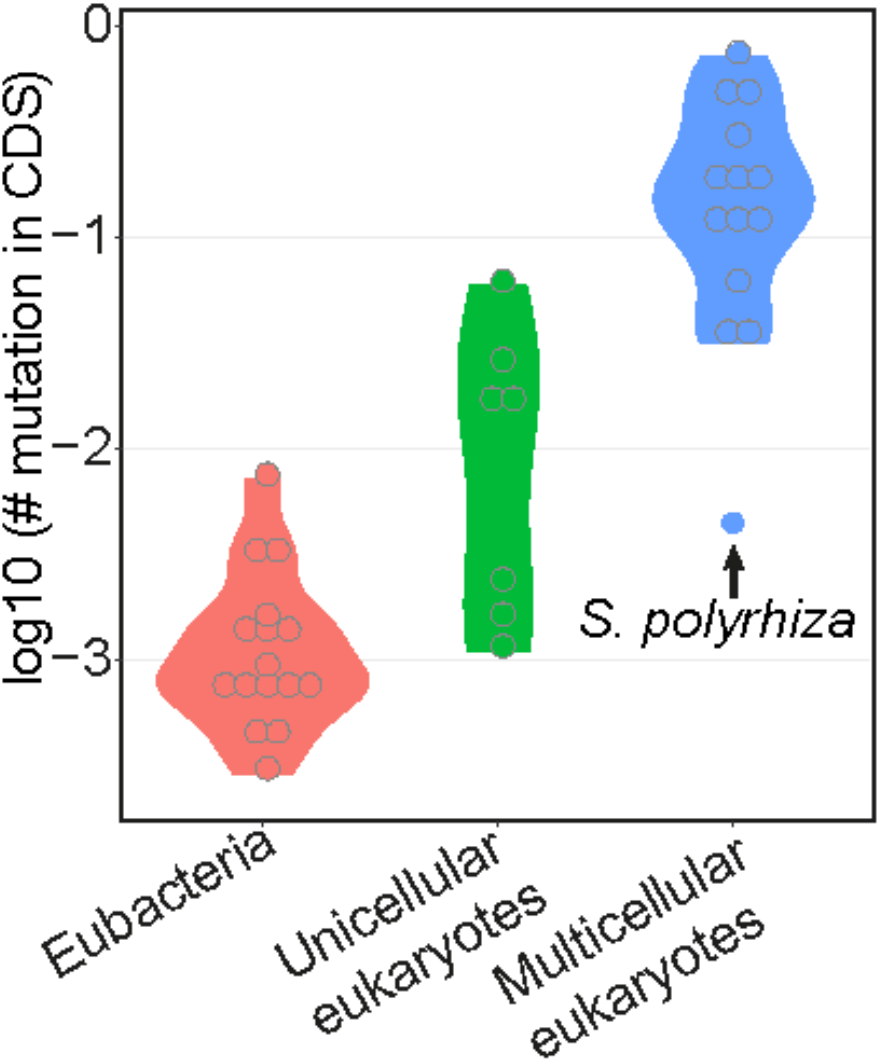
Estimated mutation rates in protein-coding regions among different organisms. The log_10_-transformed number of mutations per base pair of protein-coding genome sequences (CDS) per generation is plotted for eubacteria, unicellular eukaryotes and multicellular eukaryotes, respectively. Each gray circle indicates the estimate for one species. The arrow highlights the mutation rate in *S. polyrhiza*. Except for the mutation rate in *S. polyrhiza*, the plotted data were extracted from previous studies (Table S3).

This relatively large *N_e_* may have contributed to the evolution of a low mutation rate in *S. polyrhiza*, as selection can effectively drive down the mutation rate in populations with large *N_e_* (*8*). In addition to large *N_e_*, clonal reproduction in *S. polyrhiza* might have contributed to the evolution of a low mutation rate. In diploid species, natural selection mainly acts against deleterious homozygous variants, which appear after recombination during sexual reproduction (*22*). During the clonal phase, recessive deleterious mutations will accumulate as heterozygotes with little fitness effects. In the sexual phase these accumulated mutations can appear as homozygotes and will subsequently experience strong purifying selection. Therefore, for species such as *S. polyrhiza* that reproduce both clonally and sexually, the frequency of asexual reproduction may be negatively correlated with the mutation rate. As ~80% of all angiosperms (*23*), including many crop species (*24*), can reproduce clonally, variation in the frequency of sex may have large effects on the evolution of mutation rates in plants and contribute to variation in intraspecific genetic diversity among species.

A seemingly counter-intuitive prediction of the drift-barrier hypothesis is that populations with large *N_e_* can exhibit low genetic diversity, provided that mutation rate has evolved to be very low. The results presented here on *S. polyrhiza* support this prediction. The role of mutation rate in driving variation in genetic diversity has been largely ignored, because obtaining accurate estimates of genome-wide genetic diversity and spontaneous mutation rate in a range of organisms has been difficult in the past. Our study reveals that the low genomic diversity in *S. polyrhiza* largely stems from a low mutation rate and emphasizes that accurate estimates of mutation rates are important explaining patterns of genetic diversity within species.

## Materials and Methods

### Mutation accumulation experiments with *S. polyrhiza*

We performed a mutation accumulation (MA) experiment with *S. polyrhiza* for 20 generations. *Spirodela polyrhiza* plants were propagated under three conditions: i) indoors in the absence of UV light, ii) outdoors in the absence of natural UV light, and iii) outdoors in the presence of natural UV light. *Spirodela polyrhiza* genotype 7498 was pre-cultivated for three weeks in N-medium - which supports optimal growth (N-medium: 0.15 mM KH_2_PO_4_, 1 mM Ca(NO_3_)2 x 4 H_2_O, 8 mM KNO_3_, 5 μM H_3_BO_3_, 13 μM MnCl_2_ x 4 H_2_O, 0.4 μM Na_2_MoO_4_ x 2 H_2_O, 1 mM MgSO_4_ x 7 H_2_O, 25 μM FeNaEDTA) - in a climate chamber operating under the following conditions: 16h light, 8h dark; light supplied by vertically arranged neon tubes (OSRAM, Lumilux, cool white L36W/840) on each side; light intensity at plant height: 186±3 μmol s^−1^ m^−2^ outside polystyrene tubes and 142±3 μmol s^−1^ m^−2^ inside polystyrene tube; temperature: 28 °C constant; humidity: 41 %. The genotype 7498 originating from North Carolina (USA) was selected based on the existence of a clone-specific reference genome (*14*). A single frond (S1) was transferred to a transparent 50 ml polystyrene tube (28.5 x 95 mm, Kisker) containing 30 ml N-medium, covered with foam cap and incubated in a climate chamber under the above specified conditions. To obtain 6 MA lineages per treatment, the S1 ancestor was propagated according to the propagation scheme (Figure S2) every two to three days when daughter fronds had fully emerged from the mother frond. For the indoor MA lines, 6 lineages were consequently propagated as single descendants for 20 generations under the same conditions as described above over a period of six weeks. For the outdoor MA lines, plants were moved at the end of June 2016 into a sun-exposed field site in Jena, Germany (50°53’06.7”N 11°40’53.1’’E). The fronds were propagated in plastic beakers containing 180 ml N-medium that were fitted into the cavities of white polyvinyl chloride inserts (3 mm thickness) floating inside water-filled 10 L buckets. The buckets were surrounded with a 20 cm isolation layer of soil to avoid extreme temperature fluctuations and refilled with water to compensate for evaporating water whenever needed. To manipulate UV light, the buckets were covered with either UV transmitting (GS 2458, Sandrock, Germany) or UV blocking (UV Gallery100, Sandrock, Germany) Plexiglas plates with 1 – 3 cm distance between the bucket edge and the plates to allow air circulation. Each MA lineage was propagated in a separate bucket. After transplanting the fronds into the field, the buckets were shaded with two layers of green clear film for the first two days to allow plants to acclimate to outdoor conditions. The first green clear film layer was removed after two days, the second layer after four days. Plants were then propagated every two to four days for the following two months as single descendants for 20 generations. The MA lineages were randomized between the buckets every two weeks. The 20^th^ generation of the outdoor plants was moved back to the original growth chamber. To obtain genomic DNA for whole genome re-sequencing (WGS), a single frond of the 20^th^ generation of each of the indoor and outdoor MA lines and the ancestor, of which the roots and reproductive pockets were removed, was frozen in liquid nitrogen. All samples were stored at −80 °C until DNA extraction.

### DNA isolation and whole genome resequencing

The plant tissue was ground by vigorously shaking the Eppendorf tubes with three metal beads for 1 min in a paint shaker (Skandex S-7, Fluid Management, Sassenheim Holland) at 50 Hz. All DNA samples were isolated using the CTAB method (*25*) and their quantity and quality was analyzed on Qubit. The DNA samples from the MA experiments were sequenced on Illumina HiSeq 4000 at the Genomics Center of the Max Planck Institute for Plant Breeding Research in Cologne (Germany) with 150 bp paired-end reads. For the 68 *S. polyrhiza* genotypes, all genotypes of *Spirodela polyrhiza* (L.) Schleid. (listed in Table S1) were taken from the stock collection of the Department of Plant Physiology, University of Jena, Germany. Plants were then grown in N-medium (see details above) under a constant temperature of 28 °C and 41% humidity. Detailed information and origin of the 68 *S. polyrhiza* genotypes is listed in Table S1. The genomes of the 68 genotypes of *S. polyrhiza* were sequenced on Illumina HiSeq X Ten at BGI (Shenzhen, China) with 150 bp paired-end reads. On average, 48.2 million reads per genotype were generated.

### Short-read trimming, mapping and variant calling

For all sequenced short reads, low-quality reads and adaptor sequences were trimmed with AdapterRemoval v2.0 (*26*) with the parameters: --collapse --trimns --trimqualities – minlength 36. All of the trimmed reads were then mapped to the *S. polyrhiza* reference genome (*14*) using BWA-MEM (*27*) with default parameters. All reads with multiple mapping positions in the genome were removed and only the mapped reads were kept. PCR duplicates were removed using the “rmdup” function from SAMtools (*28*). The aligned reads were then used for variant (SNPs and small indels) calling using GATK v3.5 (*15*) following the suggestions on best practices (*29, 30*). In brief, the aligned reads around indels were re-aligned using “IndelRealigner”, and variants were called using the “UnifiedGenotyper” function with the option “-stand_call_conf 30 -stand_emit_conf 10”. The variants were then filtered with the option ““MQ0 >= 4 && ((MQ0 / (1.0 * DP)) > 0.1) & QUAL < 30.0 & QD < 5.0””. The variant clusters were further annotated as more than three variants within 50 bp using the GATK “VariantFiltration” function. Only biallelic loci were kept for downstream analysis. The synonymous and non-synonymous variants were annotated using snpEFF (version 4.3m) (*31*). Due to low sequencing coverage, three individuals from the MA experiments were removed from downstream analysis (Figure S2).

### Population genomic analysis

To analyze genetic diversity and population genomics of the 68 genotypes, additional filtering steps “-s -f “DP > 510 & DP < 10200”” were performed using vcffilter (https://github.com/vcflib/vcflib#vcffilter). Loci with missing data, variants from mitochondrial and chloroplast regions and clustered variants were removed using vcftools (*32*). The population structure among the sequenced 68 genotypes was analyzed using fastSTRUCTURE v1.0 (*33*). Multiple K values (refers to number of populations) ranging from 1 to 10 were analyzed and the value K = 4 was selected using the chooseK.py function from the fastSTRUCTURE package. The genome-wide intra-specific diversity was analyzed using Popgenome v2.2.0 (*34*) and diversity at synonymous and non-synonymous sites was analyzed using SNPGenie (*35*). Plink (*36*) was used to calculate pairwise linkage disequilibrium (LD) from the dataset; related individuals were removed and only SNPs with minor allele frequency (MAF) greater than 0.05 were kept. To model the decline of LD with physical distance, pairwise *r*^2^ between sites was used as the use of D is sensitive to small sample sizes (*37, 38*), and the decline of LD was modeled using Sved’s equation: E(*r*^2^) = (1-/(1+4 βd))+1/n, where β is the decline in LD with distance d (*39*) and 1/n accounts for small sample size (*40*). The extent of useful LD for mapping can be defined as *r*^2^ = 0.2 (*41*). In this study we use mean *r*^2^ for non-overlapping 100-bp bins to fit Sved’s equation.

### Mutation rate estimation and false-negative calculations

Accurately estimating mutation rate requires a step-wise filtering and quality checking process. The SNP filtering pipeline for the MA experiments was developed based on previous studies (*42, 43*) and iterative manual inspections of the BAM files using Integrative Genomics Viewer (IGV) (*44, 45*). 1) To reduce false positives, we only considered the mapped and properly paired reads with insertion size greater than 100 bp and less than 600 bp using bamtools (*46*). 2) We also excluded all genomic regions that were supported by fewer than nine or greater than 75 reads per sample from both variant counting and genome size calculation, as the variants from the regions that have low or high coverage are likely due to mapping errors (such as repetitive or duplicated regions). On average, 79.7% of the genomic region was kept. 3) Because spontaneous mutations should be only found in the offspring samples but not the ancestor, and the likelihood of a mutation occurring at the exact same position in multiple samples is extremely low (*u*^n^, where *u* is the mutation rate, and n refers to number of samples), any variants that appeared in more than two samples were removed. 4) Only the heterozygous variants that were supported by at least three reads for both alleles were kept. After these filtering steps, 86 variants were found (Table S7). Among these, 56 were annotated as variant clusters, likely due to mapping errors. To confirm this, we re-sequenced 28 of these variants that were located in clusters using a Sanger sequencing approach and found none of them confirmed to be true mutations. Therefore, all the variants that were classified as variant clusters were removed.

After removing all variant clusters, nine SNPs and 21 indels remained. Among the 21 indels, all of them were loss of heterozygosity in either the ancestral or offspring samples. Inspecting the alignment using the IGV showed that 19 of them were located in regions of simple sequence repeats or transposable elements, which were likely false positives. To confirm this, we selected 11 indels for Sanger sequencing and found that all of them were indeed false positives. As a result, all 21 indels were removed from the downstream analysis. Among the nine SNPs, six were point mutations (due to spontaneous mutations) and three were loss of heterozygosity (LOH) mutations (potentially due to gene conversion events). We further validated these SNPs using a Sanger-sequencing approach. Two LOH loci were very close to the gap of the genome assembly and the PCR primers could thus not be designed. We validated the remaining seven loci (six point mutations and one loss of heterozygosity). In total, four out of the six point-mutations were confirmed, and the loss of heterozygosity mutation turned out to be a false positive. The confirmed point-mutations are listed in Table 1 and were used for calculating the spontaneous mutation rate.

The relatively stringent parameters in the variant filtering process theoretically could result in a high rate of false negatives. To control this, we further estimated the false negative rate using the sequence data. We first identified all high-quality heterozygous SNP loci (30,392) from the ancestor using the same filtering parameters (coverage between nine and 75, and at least three reads to support each of the reference and the alternative allele) and compared them with the heterozygous SNPs in the offspring using a custom script. In theory, all these variants should be found in the clonally produced offspring. Thus, the number of SNPs that could not be identified from the offspring was used to estimate the highest boundary of the false negative rate from our sequencing and variant calling/filtering pipeline, as some of these cases could be a true loss of heterozygosity.

### Variant validation using Sanger sequencing

Because the total amount of DNA from a single individual was limited, the variant validation was performed using the descendants of the ancestor and offspring individuals. Specifically, at the end of the MA experiments, one individual of each line was propagated for four more generations under indoor conditions, after which the plants were frozen in liquid nitrogen for subsequent variant validation.

To validate the candidate variants, DNA was isolated as described above. PCR primers were designed based on the 500 bp flanking sequences. The PCR reactions were performed with goTaq DNA polymerase (Promega) using 30 PCR cycles with an annealing temperature of 58°C. The primer information is listed in supplemental Table S8. The PCR products were checked on a 1.5% agarose gel. The PCR products were then used for sequencing reactions using BigDye v3.1, and the products from the sequencing reactions were purified and sequenced on an ABI 3130XL sequencer.

## Acknowledgments

We thank Claudia Michel, Beatrice Arnold and Yuanyuan Song for their help in validating the variants from the MA experiments, Stefanie Schirmer for help with the MA experiment and DNA isolation, Daniel Veit for manufacturing the facilities for outdoor duckweed growth, Thomas Städler and Martin Schäfer for constructive discussions commenting on the manuscript. We are also grateful to Tobias Neumann for providing meteorological data.

## Author contributions

S. G., A. W. and M. H. performed the experiments, J. S., J. B. and S. X. performed data analysis, K. J. A. and K. S. S. contributed to the giant duckweed collections, A. W. and J. G. provided resources. M. H. and S. X. conceived and supervised the project, S. X. wrote the manuscript with input from all co-authors.

## Data and materials availability

Processed data are available in the manuscript or the supplementary materials. All raw DNA sequences obtained in this study are submitted to NCBI under Bioproject PRJNA476302.

## Supplementary figures and tables

**Figure S1.**
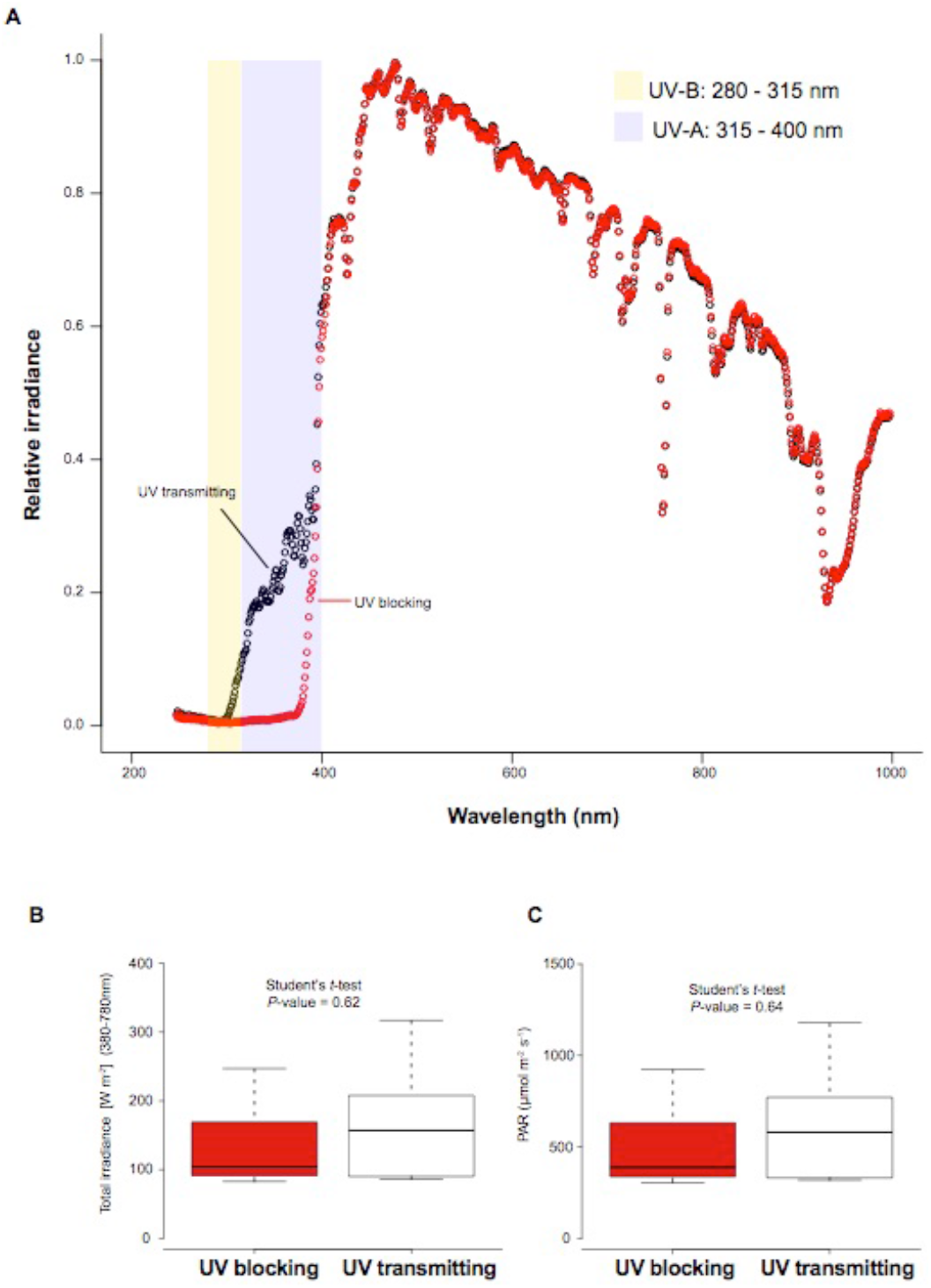
Spectral properties of UV blocking and UV transmitting plexiglass covers. A, relative irradiance (irradiance / max irradiance per plate) in UV transmitting (GS 2458, *n* = 9) and UV blocking (UV Gallery100, *n* = 9) plexiglass. B and C, total irradiance (B) and PAR (C) did not differ between UV transmitting and UV blocking plexiglass (*n* = 9). The irradiance from 250 to 1000 nm, as well as total irradiance between 380 and 780 nm and photosynthetic active radiation (PAR) between 400 and 700 nm were measured to assess the spectrum of the UV transmitting and UV blocking types of plexiglass (Sandrock, Germany) that were used in the mutation accumulation experiments.

**Figure S2.**
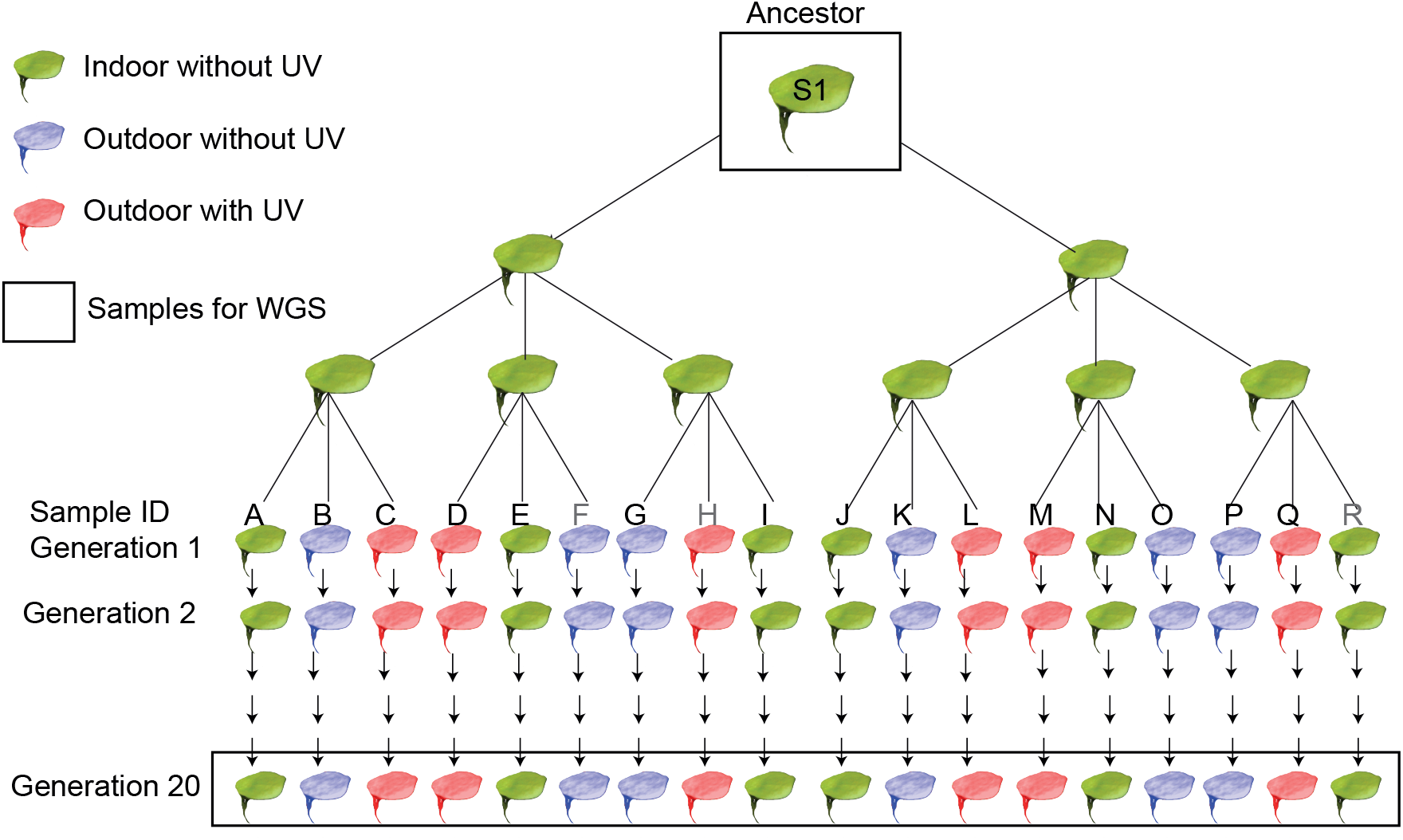
Propagating scheme of the individuals for the mutation accumulation experiments. A single ancestor was used to propagate 18 individuals, which were then propagated for 20 generations with a single-descendant approach. Among the 18 individuals collected, three individuals (F, H and R, in gray color) were not included for the data analysis due to their low sequencing depth. Each color represents a different treatment. Samples that were used for sequencing are marked with a black box (bottom row).

**Table S1.** Sample and sequencing information of 68 *S. polyrhiza* genotypes. Mapped reads refer to all uniquely mapped reads, coverage was calculated based on the nuclear genome, and accession ID refers to the registered four-digit code of each genotype. The relatively low mapping rates from some samples were mainly due to non-plant DNAs.

| Sample ID | Reads<br>sequenced<br>(million) | Mapped<br>reads<br>(million) | Mapping rate<br>(%) | Coverage<br>(X) | Accession<br>ID | Continent | Country |
| --- | --- | --- | --- | --- | --- | --- | --- |
| Sp1 | 40.1 | 19.4 | 48.47 | 16.4 | 0040 | Asia | China |
| Sp10 | 47.3 | 34.3 | 72.37 | 27.4 | 8790 | America | Canada |
| Sp100 | 49.2 | 34.2 | 69.53 | 29.7 | 5525 | Asia | Thailand |
| Sp101 | 46.5 | 32.8 | 70.44 | 26.8 | 5526 | Asia | Thailand |
| Sp105 | 55.3 | 30.1 | 54.37 | 24.9 | 10F-IP-112 | Europe | Switzerland |
| Sp106 | 52.7 | 37.1 | 70.33 | 30.7 | S5 | Europe | Switzerland |
| Sp11 | 49.3 | 33.2 | 67.38 | 28.9 | 9242 | America | Ecuador |
| Sp12 | 39.9 | 29.9 | 75.05 | 23.7 | 9503 | Asia | India |
| Sp13 | 48.6 | 38.1 | 78.34 | 32.9 | 9507 | Asia | Russia |
| Sp14 | 47.1 | 33.2 | 70.58 | 28.5 | 9509 | Europe | Germany |
| Sp15 | 41.7 | 33.4 | 80.09 | 27.5 | 9510 | Africa | Mozambique |
| Sp16 | 48.4 | 36.8 | 76.10 | 31.8 | 9636 | Asia | China |
| Sp17 | 50.0 | 38.5 | 77.15 | 32.1 | 9907 | Asia | Bangladesh |
| Sp18 | 56.7 | 44.1 | 77.76 | 38.8 | 9511 | Asia | Russia |
| Sp19 | 47.0 | 35.9 | 76.35 | 28.8 | 9925b | Asia | Bangladesh |
| Sp20 | 51.9 | 38.9 | 74.87 | 33.0 | 9625 | Europe | Albania |
| Sp21 | 47.1 | 31.1 | 65.96 | 26.3 | 9500 | Europe | Germany |
| Sp22 | 69.7 | 53.1 | 76.19 | 45.8 | 9628 | Europe | Albania |
| Sp23 | 48.8 | 35.0 | 71.78 | 29.6 | 9618 | Europe | Italy |
| Sp24 | 43.7 | 34.1 | 78.02 | 28.4 | 5523 | Asia | China |
| Sp25 | 52.2 | 37.9 | 72.57 | 31.2 | 9351 | Asia | Vietnam |
| Sp26 | 55.1 | 42.6 | 77.25 | 36.3 | 9633 | Europe | Albania |
| Sp30 | 44.3 | 29.7 | 66.94 | 26.3 | 8442 | Asia | India |
| Sp31 | 50.1 | 37.1 | 73.93 | 32.0 | 9514 | Europe | Austria |
| Sp32 | 45.1 | 35.8 | 79.34 | 30.4 | 9505 | America | Cuba |
| Sp34 | 49.2 | 33.6 | 68.29 | 28.0 | 9502 | Europe | Ireland |
| Sp35 | 45.3 | 27.5 | 60.70 | 22.9 | 0109 | Asia | China |
| Sp36 | 48.3 | 36.1 | 74.67 | 31.6 | 0092 | Asia | China |
| Sp38 | 48.3 | 36.8 | 76.29 | 32.1 | 5521 | Asia | China |
| Sp39 | 44.0 | 32.8 | 74.38 | 28.5 | 9650 | Asia | India |
| Sp4 | 44.2 | 30.1 | 68.08 | 25.1 | 7498 | America | USA |
| Sp40 | 53.5 | 36.7 | 68.47 | 32.7 | 9295 | Asia | India |
| Sp41 | 43.7 | 35.7 | 81.76 | 30.4 | 9609 | Europe | Poland |
| Sp42 | 52.2 | 35.8 | 68.53 | 29.4 | 9629 | Europe | Albania |
| Sp43 | 49.4 | 28.3 | 57.39 | 24.4 | 9497 | Asia | India |
| Sp45 | 45.1 | 36.1 | 79.98 | 30.7 | 9512 | Asia | Russia |
| Sp46 | 50.6 | 39.8 | 78.60 | 33.1 | 9504 | Asia | India |
| Sp47 | 48.1 | 26.1 | 54.29 | 21.6 | 9613 | Europe | Poland |
| Sp48 | 45.7 | 28.2 | 61.74 | 23.6 | 9305 | Asia | India |
| Sp5 | 40.4 | 25.0 | 61.85 | 21.6 | 7551 | Asia | Australia |
| Sp50 | 44.1 | 35.9 | 81.51 | 29.7 | 0090 | Asia | China |
| Sp51 | 48.1 | 26.6 | 55.27 | 22.0 | 9316 | Asia | India |
| Sp53 | 41.6 | 28.1 | 67.65 | 22.7 | 9290 | Asia | India |
| Sp54 | 42.1 | 25.6 | 60.83 | 21.1 | 8787 | Asia | Nepal |
| Sp55 | 45.5 | 34.0 | 74.55 | 29.1 | 9610 | Europe | Poland |
| Sp56 | 45.9 | 26.3 | 57.33 | 21.9 | 9256 | Europe | Finland |
| Sp57 | 48.0 | 34.2 | 71.16 | 28.3 | 9333 | Asia | China |
| Sp58 | 43.3 | 33.8 | 77.92 | 27.2 | 9560 | Europe | Hungary |
| Sp59 | 44.9 | 35.2 | 78.26 | 29.4 | 9506 | Asia | India |
| Sp61 | 43.7 | 32.0 | 73.09 | 26.8 | 0225 | Asia | China |
| Sp62 | 46.0 | 19.6 | 42.54 | 16.5 | 9513 | Europe | Czech Republic |
| Sp63 | 44.2 | 26.5 | 59.97 | 22.3 | 0013 | Asia | Vietnam |
| Sp65 | 53.6 | 41.2 | 76.77 | 34.6 | 9413 | Europe | Italy |
| Sp66 | 54.8 | 41.1 | 74.98 | 30.7 | 9622 | Europe | Germany |
| Sp7 | 44.1 | 32.9 | 74.50 | 27.7 | 7674 | Asia | Nepal |
| Sp71 | 49.5 | 27.4 | 55.26 | 24.0 | 9608 | Europe | Poland |
| Sp72 | 50.4 | 29.0 | 57.59 | 23.4 | 5522 | Asia | China |
| Sp77 | 55.2 | 43.1 | 78.14 | 37.3 | 9508 | Europe | Poland |
| Sp78 | 47.1 | 33.6 | 71.41 | 29.3 | 9607 | Europe | Switzerland |
| Sp79 | 49.2 | 35.4 | 71.94 | 30.9 | 5513 | Europe | Germany |
| Sp8 | 43.7 | 34.6 | 79.04 | 28.9 | 8683 | Africa | Kenya |
| Sp82 | 44.1 | 33.6 | 76.27 | 28.7 | 9649 | Asia | India |
| Sp85 | 50.0 | 38.0 | 76.06 | 34.0 | 9192 | America | Colombia |
| Sp87 | 55.6 | 39.6 | 71.28 | 34.9 | 9657 | America | Canada |
| Sp89 | 58.3 | 41.6 | 71.30 | 37.2 | 0192 | Asia | China |
| Sp9 | 40.8 | 29.1 | 71.33 | 25.0 | 8756 | Africa | Ethiopia |
| Sp93 | 52.5 | 41.8 | 79.48 | 33.7 | 9346 | Europe | Switzerland |
| Sp99 | 54.9 | 37.3 | 67.93 | 31.8 | 5524 | Asia | Thailand |

**Table S2.** Summary statistics of population genomics in *S. polyrhiza*. π referes to the estimated pairwise nucleotide diversity. π_a_ refers to the nucleotide diversity at non-synonymous sites and π_s_ refers to the nucleotide diversity at synonymous sites.

| Summary statistics | Genome-wide $\pi$ | $\pi$ in coding regions | $\pi$ in noncoding regions | Nonsyn. variants | Syn. variants | $\pi_a$ | $\pi_s$ | $\pi_a/\pi_s$ |
| --- | --- | --- | --- | --- | --- | --- | --- | --- |
| All populations combined | 0.0013 | 0.00051 | 0.0014 | 14,191 | 8,865 | 0.00038 | 0.00086 | 0.44 |

**Table S3.** Summary of mutation rate and effective population size (*N_e_*) estimates. Data is obtained from Lynch et al. 2016 (*47*), with a few updates (*48*). Mutation rate (*μ*) is listed as per generation per site. NA: not available. CDS: protein coding sequence. The estimated π from neutral sites is calculated as D × *N_e_* × *μ*, where D refers to 4 (diploid species) or 2 (haploid species).

| Species | Group | $\mu$ | $N_e$ | Genome size (Mb) | CDS size (Mb) | Mutations per CDS per generation | Estimated $\pi$ from neutral sites |
| --- | --- | --- | --- | --- | --- | --- | --- |
| *Apis mellifera* | Multicellular eukaryotes | 6.80E-09 | NA | 262.0 | 29.0 | 0.197 | NA |
| *Arabidopsis thaliana* | Multicellular eukaryotes | 6.95E-09 | 2.9E+05 | 119.7 | 42.1 | 0.292 | 0.00806 |
| *Caenorhabditis briggsae* | Multicellular eukaryotes | 1.33E-09 | 2.7E+05 | 104.0 | 24.1 | 0.032 | 0.00144 |
| *Caenorhabditis elegans* | Multicellular eukaryotes | 1.45E-09 | 5.4E+05 | 100.3 | 25.0 | 0.036 | 0.00313 |
| *Daphnia pulex* | Multicellular eukaryotes | 5.69E-09 | 8.3E+05 | 250.0 | 30.2 | 0.172 | 0.01889 |
| *Drosophila melanogaster* | Multicellular eukaryotes | 5.17E-09 | 8.6E+05 | 168.7 | 23.2 | 0.120 | 0.01778 |
| *Heliconius melpomene* | Multicellular eukaryotes | 2.90E-09 | 2.1E+06 | 273.8 | 39.1 | 0.113 | 0.02436 |
| *Homo sapiens* | Multicellular eukaryotes | 1.35E-08 | 2.1E+04 | 3300.0 | 36.5 | 0.493 | 0.00113 |
| *Mus musculus* | Multicellular eukaryotes | 5.40E-09 | 1.8E+05 | 2717.0 | 35.5 | 0.192 | 0.00389 |
| *Oryza sativa* | Multicellular eukaryotes | 7.10E-09 | 5.3E+04 | 389.0 | 101.3 | 0.719 | 0.00151 |
| *Pan troglodytes* | Multicellular eukaryotes | 1.20E-08 | 2.9E+04 | 3524.0 | 37.2 | 0.446 | 0.00139 |
| *Pristionchus pacificus* | Multicellular eukaryotes | 2.00E-09 | 1.8E+06 | 169.7 | 29.7 | 0.059 | 0.01440 |
| *Clupea harengus* | Multicellular eukaryotes | 2.00E-09 | 4.0E+05 | 850.0 | 57.6 | 0.115 | 0.00320 |
| *Spirodela polyrhiza* | **Multicellular eukaryotes** | **2.38E-10** | 9.0E+05 | **158.0** | **17.4** | **0.004** | 0.00086 |
| *Chlamydomonas reinhardtii* | Unicellular eukaryotes | 3.80E-10 | 4.3E+07 | 111.1 | 39.2 | 0.015 | 0.03268 |
| *Neurospora crassa* | Unicellular eukaryotes | 4.10E-09 | 1.8E+06 | 38.6 | 14.5 | 0.059 | 0.01476 |
| *Paramecium tetraurelia* | Unicellular eukaryotes | 1.94E-11 | 1.0E+08 | 72.1 | 56.8 | 0.001 | 0.00776 |
| *Plasmodium falciparum* | Unicellular eukaryotes | 2.08E-09 | 3.5E+05 | 22.9 | 12.1 | 0.025 | 0.00146 |
| *Saccharomyces cerevisiae* | Unicellular eukaryotes | 2.63E-10 | 7.8E+06 | 12.5 | 8.7 | 0.002 | 0.00410 |
| *Schizosaccharomyces pombe* | Unicellular eukaryotes | 2.17E-10 | 1.4E+07 | 19.6 | 7.2 | 0.002 | 0.00608 |
| *Trypanosoma brucei* | Unicellular eukaryotes | 1.38E-09 | 5.3E+06 | 26.1 | 13.2 | 0.018 | 0.02926 |
| *Agrobacterium tumefaciens* | Eubacteria | 2.92E-10 | 3.4E+08 | 5.7 | 5.0 | 0.001 | 0.19856 |
| *Bacillus subtilis* | Eubacteria | 3.35E-10 | 6.1E+07 | 4.3 | 3.6 | 0.001 | 0.04087 |
| *Burkholderia cenocepacia* | Eubacteria | 1.33E-10 | 2.5E+08 | 7.7 | 6.7 | 0.001 | 0.06650 |
| *Deinococcus radiodurans* | Eubacteria | 4.99E-10 | NA | 3.3 | 2.9 | 0.001 | NA |
| *Escherichia coli* | Eubacteria | 2.00E-10 | 1.8E+08 | 4.6 | 3.9 | 0.001 | 0.072 |
| *Helicobacter pylori* | Eubacteria | 1.90E-09 | 4.0E+07 | 1.7 | 1.5 | 0.003 | 0.152 |
| *Mesoplasma florum* | Eubacteria | 9.78E-09 | 1.1E+06 | 0.8 | 0.7 | 0.007 | 0.022 |
| *Mycobacterium smegmatis* | Eubacteria | 5.27E-10 | NA | 7.0 | 6.5 | 0.003 | NA |
| *Mycobacterium tuberculosis* | Eubacteria | 1.95E-10 | NA | 4.4 | 4.0 | 0.001 | NA |
| *Pseudomonas aeruginosa* | Eubacteria | 7.92E-11 | 2.1E+08 | 6.5 | 5.9 | 0.000 | 0.0333 |
| *Salmonella enterica* | Eubacteria | 1.74E-10 | 3.5E+08 | 4.9 | 4.0 | 0.001 | 0.1218 |
| *Salmonella typhimurium* | Eubacteria | 1.52E-10 | NA | 4.9 | 4.3 | 0.001 | NA |
| *Staphylococcus epidermidis* | Eubacteria | 7.40E-10 | 3.5E+07 | 2.6 | 2.1 | 0.002 | 0.0518 |
| *Thermus thermophilus* | Eubacteria | 1.38E-10 | 2.3E+08 | 2.1 | 2.1 | 0.000 | 0.0635 |
| *Vibrio cholerae* | Eubacteria | 1.15E-10 | 4.8E+08 | 3.9 | 3.4 | 0.000 | 0.1104 |
| *Vibrio fischeri* | Eubacteria | 2.08E-10 | NA | 4.3 | 3.7 | 0.001 | NA |

**Table S4.** Pairwise Fst between four population groups. The number of sequenced individuals is listed in bracket.

| **Population** | India (13) | SE Asia (18) | Europe (27) | America (10) |
| --- | --- | --- | --- | --- |
| India (13) | / | / | / | / |
| SE Asia (18) | 0.47 | / | / | / |
| Europe (27) | 0.65 | 0.35 | / | / |
| America (10) | 0.82 | 0.67 | 0.79 | / |

**Table S5.** Summary of nucleotide diversity (ND) in *S. polyrhiza*. The summary statistics of nucleotide diversity are shown separately for each chromosome. Ti/Tv: Transition to Transversion ratio. Fst is calculated among all four populations. π: average pairwise nucleotide diversity from all sites.

| Chr ID | Size (bp) | Ti/<br>Tv | Fst | $\pi$ in<br>India | $\pi$ in SE<br>Asia | $\pi$ in<br>Europe | $\pi$ in<br>America |
| --- | --- | --- | --- | --- | --- | --- | --- |
| ChrS01 | 11466534 | 2.16 | 0.67 | 0.00026 | 0.00050 | 0.00030 | 0.00018 |
| ChrS02 | 8941172 | 2.21 | 0.67 | 0.00026 | 0.00056 | 0.00033 | 0.00015 |
| ChrS03 | 8796147 | 2.23 | 0.73 | 0.00026 | 0.00061 | 0.00034 | 0.00014 |
| ChrS04 | 8491500 | 2.18 | 0.69 | 0.00025 | 0.00051 | 0.00035 | 0.00011 |
| ChrS05 | 8389602 | 2.28 | 0.66 | 0.00042 | 0.00062 | 0.00039 | 0.00021 |
| ChrS06 | 8130874 | 2.24 | 0.71 | 0.00023 | 0.00058 | 0.00033 | 0.00018 |
| ChrS07 | 8107549 | 2.30 | 0.70 | 0.00041 | 0.00060 | 0.00024 | 0.00017 |
| ChrS08 | 7340019 | 2.09 | 0.68 | 0.00028 | 0.00049 | 0.00027 | 0.00014 |
| ChrS09 | 7208038 | 2.19 | 0.68 | 0.00033 | 0.00055 | 0.00035 | 0.00017 |
| ChrS10 | 7041313 | 2.25 | 0.66 | 0.00036 | 0.00057 | 0.00042 | 0.00022 |
| ChrS11 | 6552830 | 2.20 | 0.71 | 0.00026 | 0.00054 | 0.00029 | 0.00017 |
| ChrS12 | 5946178 | 2.18 | 0.62 | 0.00037 | 0.00067 | 0.00043 | 0.00024 |
| ChrS13 | 5476630 | 2.20 | 0.70 | 0.00024 | 0.00047 | 0.00036 | 0.00016 |
| ChrS14 | 5103705 | 2.19 | 0.63 | 0.00034 | 0.00061 | 0.00039 | 0.00025 |
| ChrS15 | 4726429 | 2.38 | 0.65 | 0.00034 | 0.00061 | 0.00038 | 0.00019 |
| ChrS16 | 4623610 | 2.40 | 0.72 | 0.00027 | 0.00070 | 0.00029 | 0.00020 |
| ChrS17 | 4564609 | 2.30 | 0.67 | 0.00038 | 0.00069 | 0.00037 | 0.00016 |
| ChrS18 | 4370269 | 2.31 | 0.71 | 0.00030 | 0.00056 | 0.00040 | 0.00013 |
| ChrS19 | 3727809 | 2.26 | 0.65 | 0.00038 | 0.00058 | 0.00039 | 0.00016 |
| ChrS20 | 3541257 | 2.11 | 0.69 | 0.00030 | 0.00062 | 0.00029 | 0.00020 |

**Table S6.** Summary of the information of the sequencing coverage for the mutation accumulation experiments. The coverage was calculated based on all properly mapped reads after removing the PCR duplicates. All sample IDs refer to the samples showed in Figure S2, except sample V, which refers to the ancestor (labeled as S1 in Figure S2).

| Sample ID | Coverage (X) |
| --- | --- |
| A | 29 |
| B | 27 |
| C | 30 |
| D | 36 |
| E | 22 |
| G | 33 |
| I | 34 |
| J | 32 |
| K | 20 |
| L | 24 |
| M | 34 |
| N | 28 |
| O | 23 |
| P | 27 |
| Q | 23 |
| V | 29 |

**Table S7.**
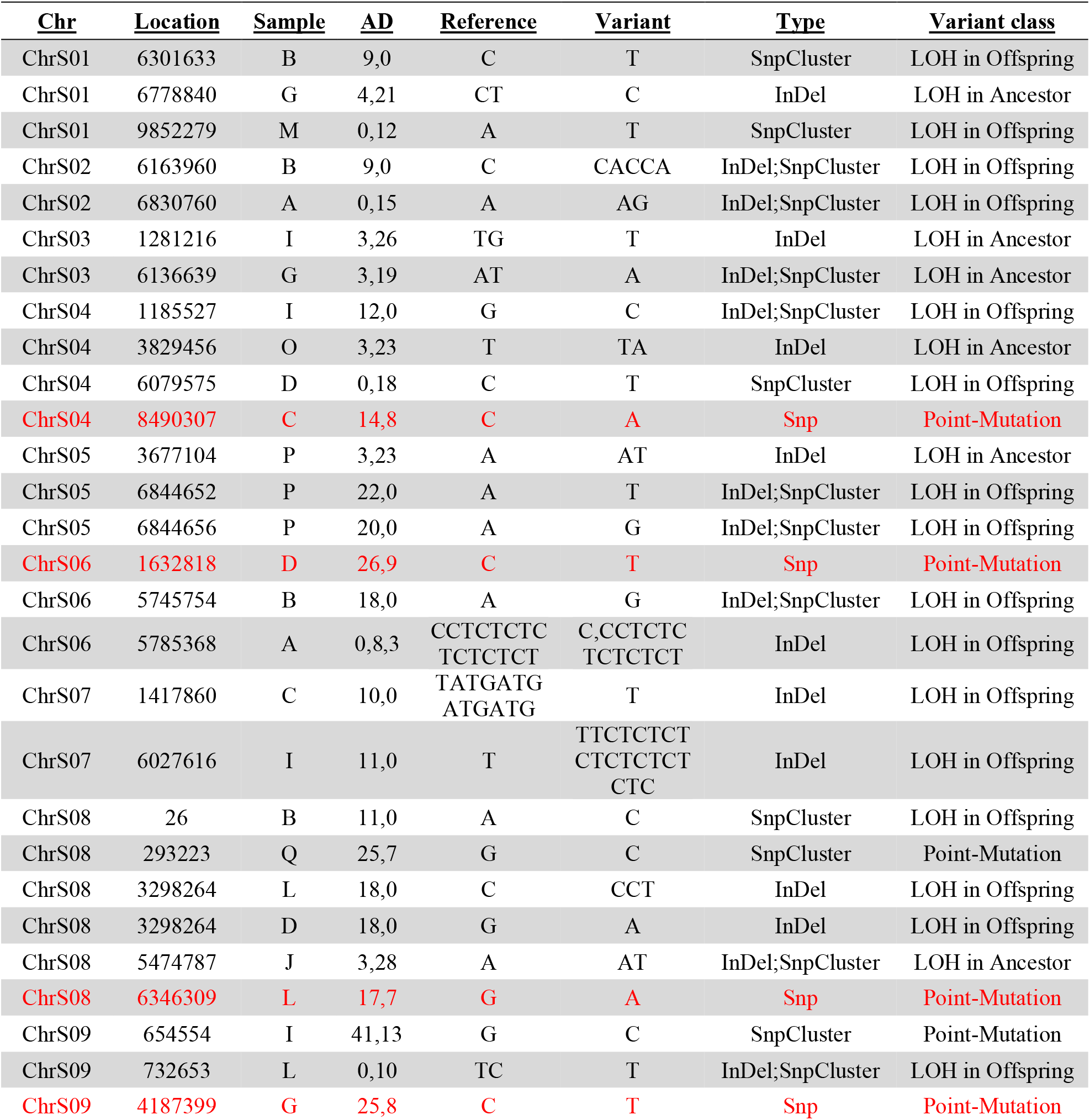
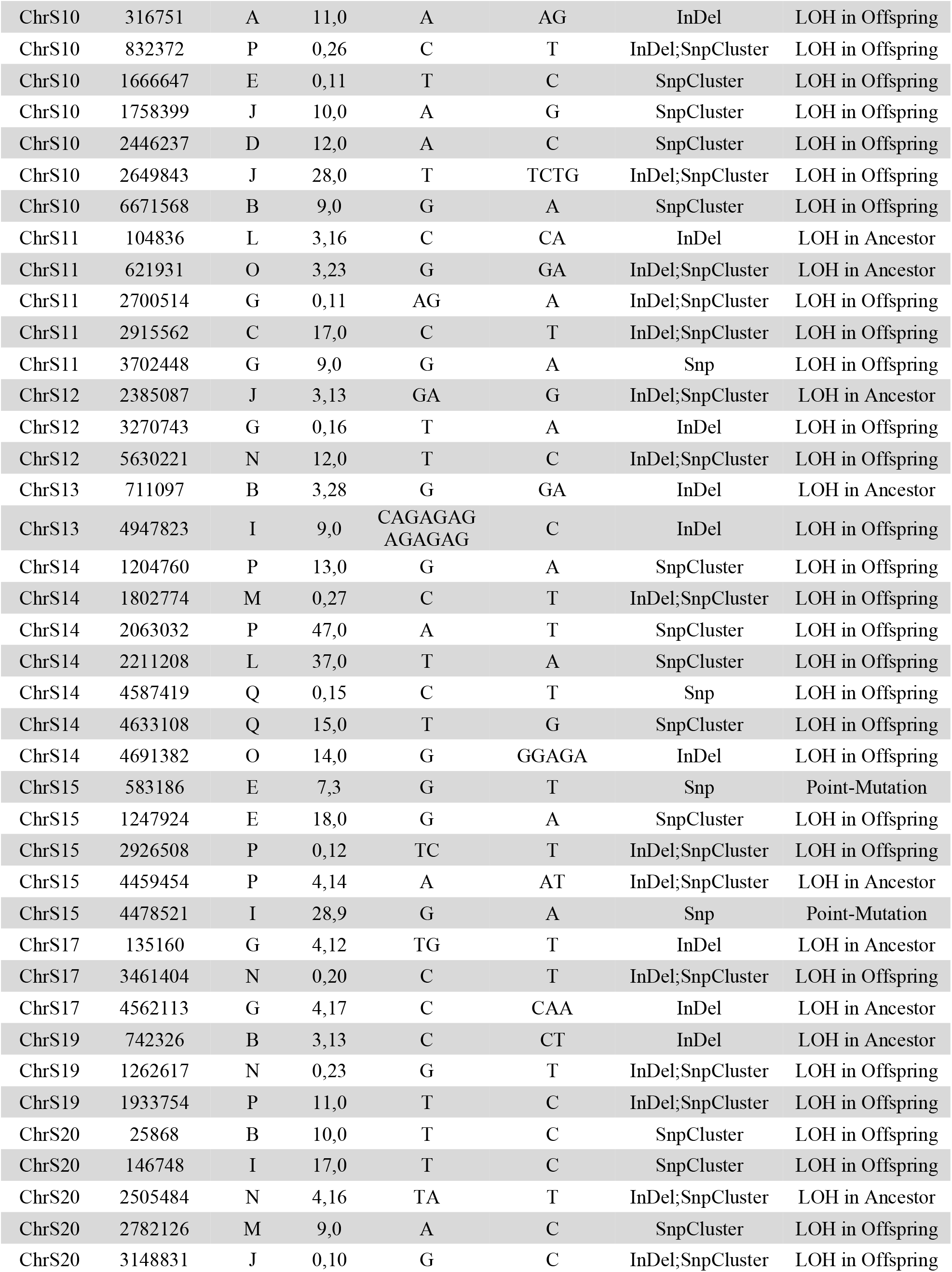
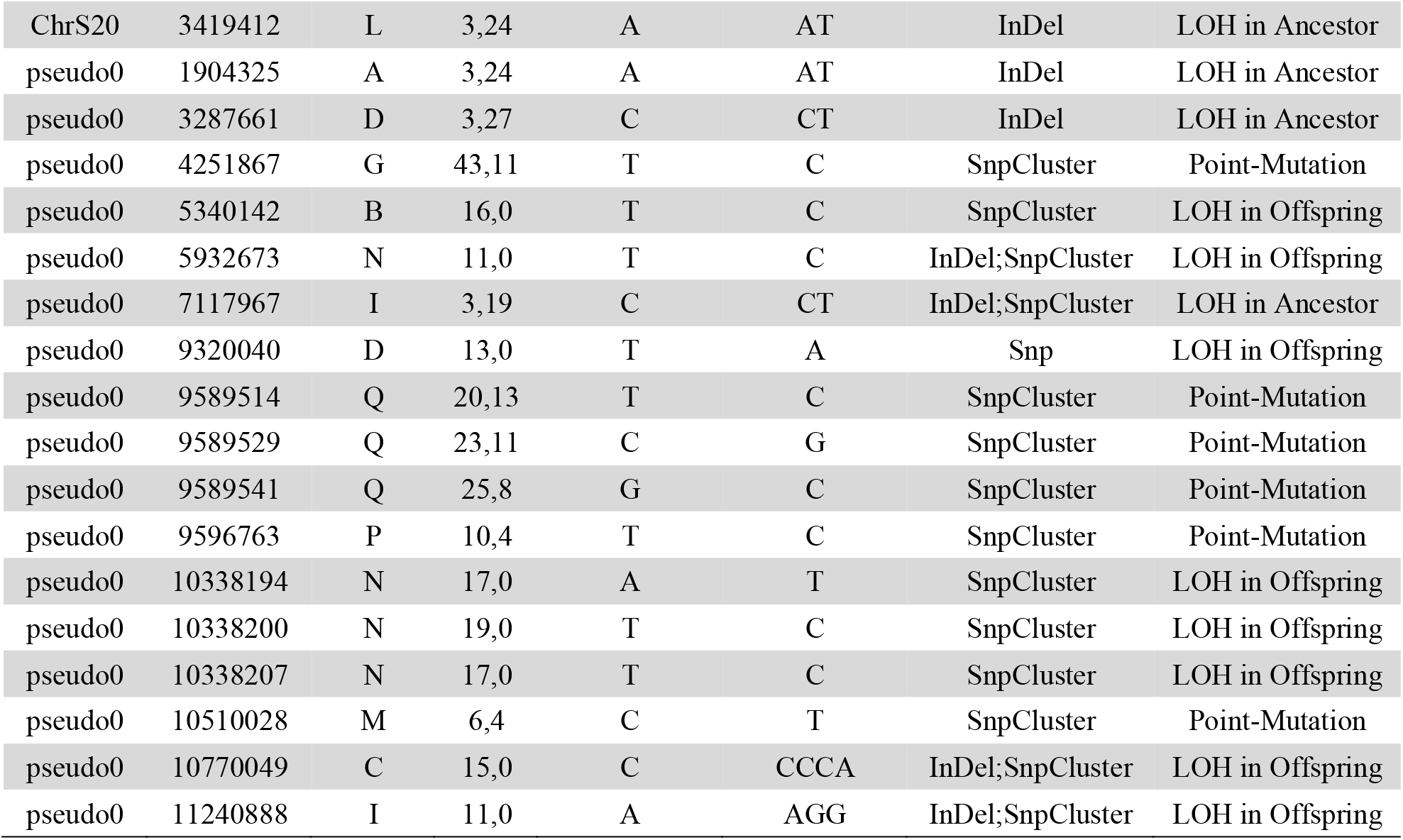
Detailed information for all putative MA variants. Most of the putative variants were loss-of-heterozygosity (LOH) mutations and located in clusters, likely due to false positives. The variants that were validated using Sanger sequencing are highlighted in red, which are all located in non-coding region. AD refers to the number of reads supporting reference and alternative alleles, respectively.

**Table S8.**
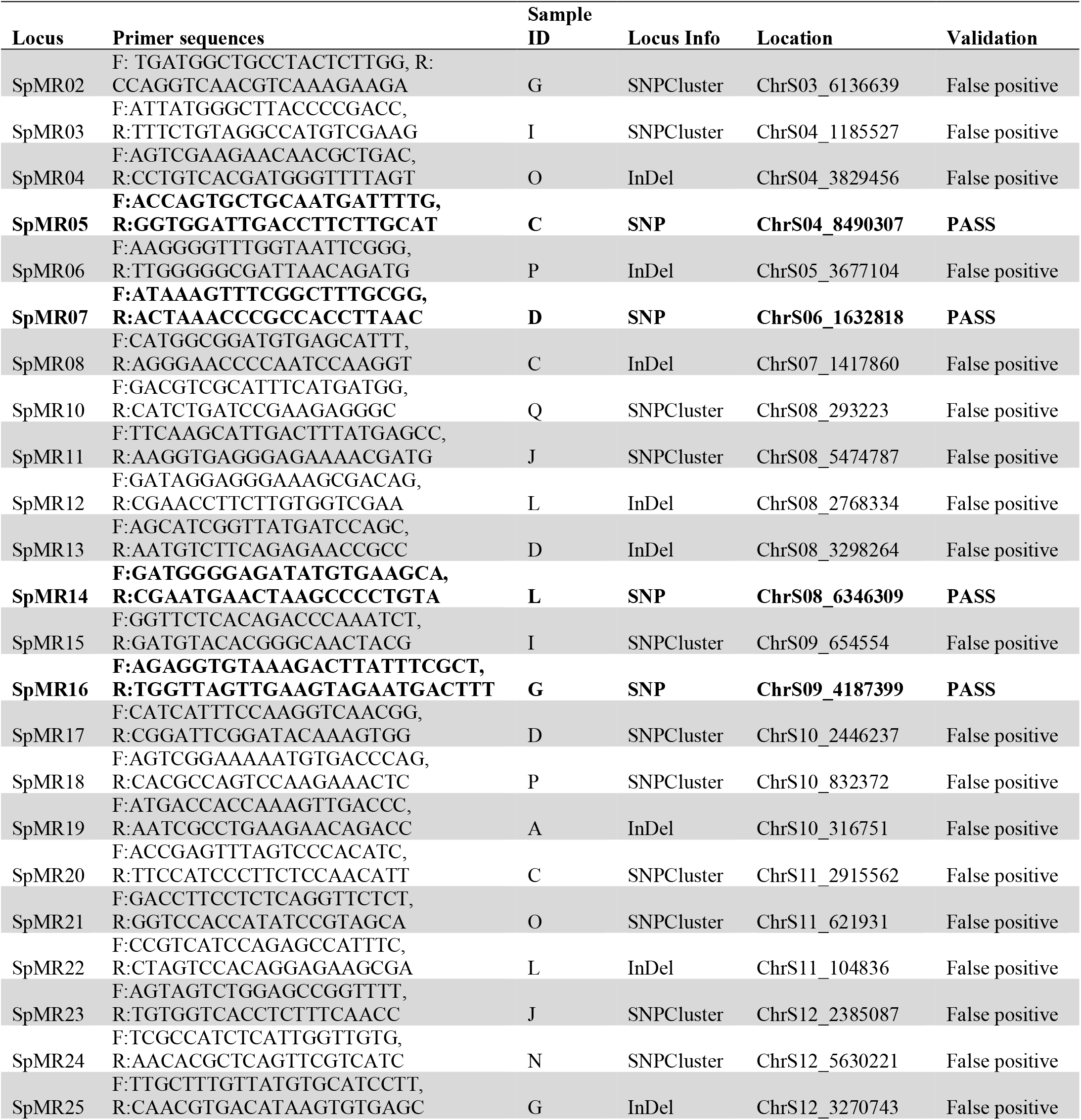
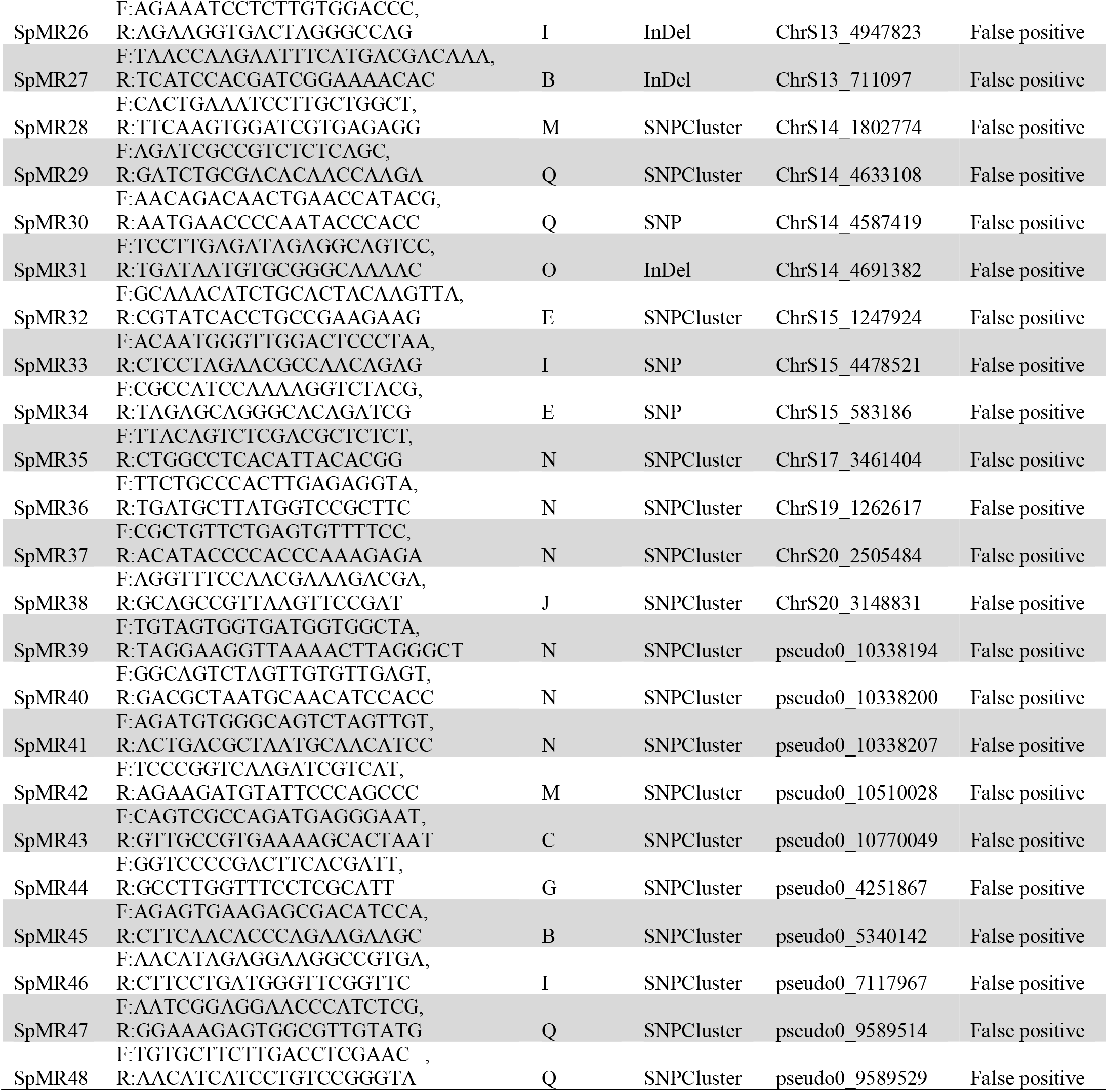
Primer information for validating the variants. All primers that were used for validating the candidate variants are shown. Primer sequence information is shown in forward (F) and reverse (R). The validation results are indicated in bold text.

**External Dataset 1. Annotation of SNPs at the gene level**. The total number of SNPs that were found in each gene is listed according to the predicted effects.

**External Dataset 2. Climate and light spectrum information in Jena (Germany), at the place where the outdoor mutation accumulation experiments were performed**. Ambient temperature, global radiation, PAR radiation and the UV spectrum are shown hourly.

